# Comparative transcriptomic analysis reveals signatures of selection for orb-weaving behavior in spiders

**DOI:** 10.64898/2026.03.30.715290

**Authors:** Calvin Runnels, Jeremiah Miller, Andrew Gordus

## Abstract

Spiders (Araneae) are responsible for one of the most captivating and intricate examples of animal architecture in the natural world: the web. But only certain groups of spiders weave the familiar spiral-shaped orb web, and the evolutionary origin of orb-weaving has left arachnologists tangled in a debate for the past decade. Since phylogenetic studies rejected the long-held belief that orb-weavers were monophyletic, two competing hypotheses have emerged: that the cribellate (dry) and ecribellate (sticky) orb webs arose independently and convergently, or that the ability to weave the orb originated with web-spinning common ancestors of all extant orb-weavers and was subsequently lost in many non-orb-weaving descendants. Attempts to reconstruct the ancestral state of the orb web have reached conflicting conclusions as a result of disagreements about the species phylogeny and the definition of orb-weaving. As a potential solution to this phylogenetic impasse, we tested orthologous genes from across the genomes of 98 species of spiders for evidence of both convergent positive selection and relaxed selection corresponding to the orb-weaving phenotype, the patterns that we would expect to be caused by each of the competing hypotheses on the origins of the orb. Using a permutation-based approach, we also compared the odds of gene loss and duplication between orb-weavers and non-orb-weaving spiders and identified genes whose copy number differ significantly between the two phenotypic groups. Through these analyses, we integrate the evolutionary history and genetic basis of orb web-associated traits, providing unique insights into the emergence of complex behavior.

## Introduction

Orb-weaving, the behavioral program used by some spiders to generate the iconic wheel-shaped orb web, is an attractive model for studying the genetic and neuronal basis of complex, innate behaviors (Corver et al. 2021; Quesada-Hidalgo et al. 2021; Artiushin et al. 2024). Although all spiders produce multiple different types of silk, not all lineages of spiders construct orb webs; some are sheet weavers, cob weavers, funnel weavers, and builders of various other types of web (Blamires et al. 2017). Some spiders make no capture web at all, instead using specialized silk types to protect their eggs, line their burrows, fish for moths, or even scuba dive (Yeargan 1988; Seymour and Hetz 2011; Uchman et al. 2018). It was long assumed that, because such a meticulous and elaborate behavior as orb-weaving could presumably have evolved no more than once, all the orb-weavers must belong to a single clade of the spider phylogeny, termed Orbiculariae (Griswold et al. 1998; Hormiga and Griswold 2014; Eberhard and Barrantes 2015). However, more recent phylogenetic studies using modern molecular tools have shown that the orb-weavers are not monophyletic. Instead, a vast evolutionary gulf appears to separate the most distantly related orb-weaving families (Bond et al. 2014; Fernández et al. 2014). The most recent branch of Araneae to contain all the orb-weavers, along with many non-orb-weaving relatives, is Entelegynae, a ∼200-million-year-old clade named for an innovation in their fertilization system (Kulkarni et al. 2023). The most speciose and well known group of orb-weaving spiders, the araneids, make the largest and strongest orb webs, and they are also the evolutionarily youngest family of spiders with the trait (Garb et al. 2019). The other two major groups are the long-jawed orb-weavers, or tetragnathids, and the cribellate orb-weavers, named for their cribellum, a loom-like anatomical structure composed of a fused row of spinnerets which they use to generate their form of capture silk. Unlike the sticky, droplet-forming aggregate glue of the araneoid orb-weavers, cribellate silk has a woolly texture that achieves a mechanical form of adhesion (Correa-Garhwal et al. 2022). Although the silk-spinning anatomy and the chemistry of the capture silk differs between distantly related orb-weavers, the silk types involved in building all other parts of the web are conserved across all orb-weavers (Tarakanova and Buehler 2012; Wolff et al. 2024). This combination of apparently divergent and conserved behaviors, anatomical features, and silk proteins has led to a fierce debate as to whether the orb web is indeed ancestral to both the cribellate and ecribellate orb-weavers, or whether it might, in fact, have evolved more than once (Dimitrov et al. 2011; Kallal et al. 2021).

Previous studies have used ancestral trait reconstruction to investigate the evolutionary history of orb-weaving (Blackledge et al. 2009; Fernández et al. 2018; Coddington et al. 2019). Such analyses infer relative probabilities of gaining or losing traits at branch points in a tree based on phylogenetic relationships. However, the higher-level relationships among spider species are in some cases still a matter of controversy because of disagreements among phylogenies derived from different data sources (Kulkarni et al. 2021). The use of different phylogenies to infer ancestral characters, along with debates on how those characters should be defined, has led to conflicting results. Furthermore, analyses that use ancestral state reconstruction do not associate these traits with specific genes or loci, but only with species-level relationships. Ultimately, the orb-weaving behavior is the result of morphological and developmental differences that are encoded by genes. If the trait is ancestral, we might expect a core set of relevant genetic elements to be conserved in orb-weavers but less well conserved in extant non-orb-weaving descendants of that same common ancestor. If the trait has evolved separately and convergently, the shared constraints on development and the genome could favor changes in the same sets of genes, albeit harboring potentially distinct mutations (Treaster et al. 2021). Thus, we might expect all orb-weavers to share complements of genes with adaptations relevant to the behavior, even if those adaptations arose independently in evolution.

In this study, we take advantage of the rich array of publicly available genetic data and powerful evolutionary hypothesis testing tools to perform an unbiased search for genes involved in orb-weaving. We tested orthologous gene sets from 98 species of entelegyne spiders for patterns of selection that correspond to either hypothesis: relaxed selection in non-orb-weavers relative to their orb-weaving relatives, as expected under the ancestral orb-weaving model, and convergent positive selection in the orb-weavers, as expected under the independent origins model. Through these approaches, we identified 1,637 orthologous genes with patterns of selection that map onto the distribution of the orb-weaving trait across the spider family tree. To address the potential contributions of gene loss and duplication, which might also have played an important role in the evolution of orb-weaving but cannot be assessed using tools which rely on comparison of substitution rates between sequences, we also developed a method for identifying genes with patterns of loss or gain that correlate with orb-weaving. We identified 683 orthologous genes whose odds of having been lost or duplicated among entelegyne spiders are significantly different between orb-weavers and non-orb-weavers. Using the annotated genome of the model cribellate orb-weaver *Uloborus diversus* as a representative example (Miller et al. 2023), we found that the putative selected genes from these analyses represent known silk gland genes as well as predicted nervous system, body planning, and sex-related genes.

## Results

### Orthologous gene discovery in entelegyne spiders

We developed a pipeline to perform comparative analyses of thousands of protein-coding genes from a phylogeny of nearly one hundred spider species (**Fig. 1a**). To ensure comprehensive representation across spiders, we compiled transcriptomes of 573 spider species from NCBI (**Supplementary Table S1**). We filtered these transcriptomes for completeness and redundancy, leaving a substantial dataset of 102 spider species used for orthologous gene discovery (**Fig. 1b**). We selected the fruit fly *Drosophila melanogaster* as a non-spider outgroup species. The resulting phylogeny includes species from across the spider tree of life, including mygalomorph and synspermiate spiders, cob weavers, sheet weavers, non-web-building spiders, and orb weavers of the three major lineages: the cribellate orb-weavers, the ecribellate tetragnathids, and the ecribellate araneids (**Figure 1c–f**). We used OrthoFinder (Emms and Kelly 2019; Emms et al. 2025) to identify 46,218 sets of orthologous genes, or orthogroups, across these 103 species. OrthoFinder further resolved these orthogroups into phylogenetic hierarchical orthogroups (HOGs). Within the subtree containing all the entelegyne spiders, and thus all extant orb-weavers, OrthoFinder recovered 107,987 HOGs, which range from species-specific gene families to genes with at least one ortholog in all 98 entelegyne spider species (**Supplementary Table S2**). OrthoFinder did not find any single-copy, full-occupancy orthogroups, likely due to widespread paralogy in spider genomes resulting from an ancient whole-genome duplication in the ancestor of the arachnopulmonates (Schwager et al. 2017; Li et al. 2026).

**Figure 1:**
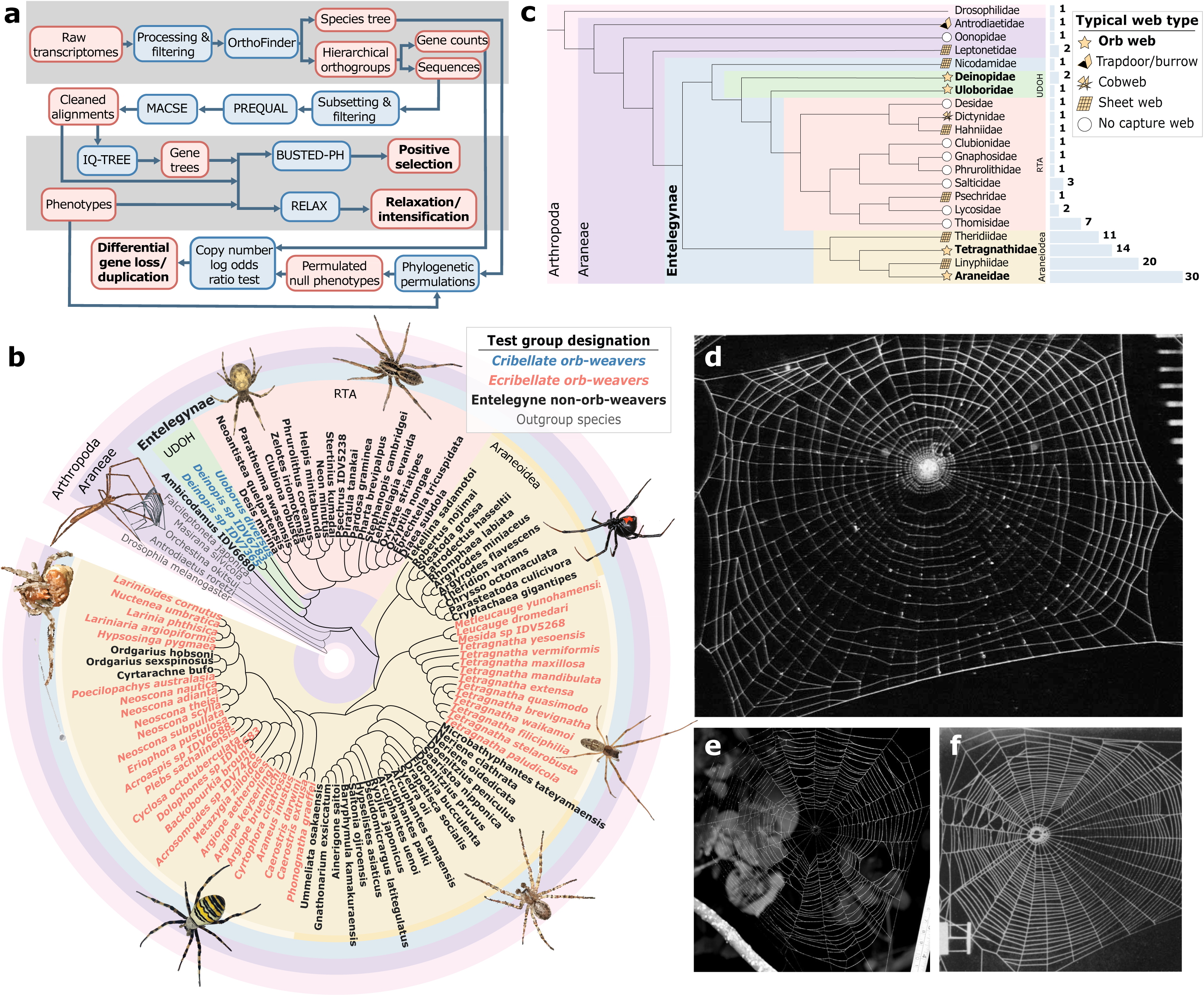
Overall pipeline and phylogenetic relationships of species included. **(a)** Flowchart of the computational pipeline. **(b)** Species tree of the 103 species used in the orthology search, including one non-arachnid outgroup species, four non-entelegyne species, three cribellate orb-weavers, and 41 ecribellate orb-weavers (14 tetragnathids and 27 araneids) (Hedin 2008; Angelus 2011; Gallagher 2013; Gallagher 2017; Hodnett 2021; Schoenmakers 2021; Anon 2022). See Methods, Orb-weaving phenotype designation. **(c)** Cladogram showing family-level relationships among analyzed species and typical web types (or lack thereof) produced by each family. Blue bars indicate counts of species per family included in the analysis. **(d)** Web of adult female *U. diversus*, a cribellate orb-weaver (Eberhard 1972). **(e)** Web of a tetragnathid (ecribellate) orb-weaver, species unspecified (Nasir 2016). **(f)** Web of an araneid (ecribellate) orb-weaver, *Neoscona vertebrata* (Baum and Witt 1960).

### Testing phylogenetic models of selection shaping orb-weaving

To mitigate the effect of missing data in the alignments, we filtered the entelegyne HOGs for a minimum occupancy of 75 (orthogroups that contain at least one representative gene from at least 75 of the 98 of species) and for the presence of at least one gene from *U. diversus*, to enable downstream functional analysis and eventually experimental validation. For each orthogroup, we generated multiple sequence alignments of the coding sequences and inferred gene trees from these alignments. We tested gene trees and alignments for evidence of positive or relaxed selection using the HyPhy tests RELAX and BUSTED (Murrell et al. 2015; Wertheim et al. 2015; Kosakovsky Pond et al. 2020). These maximum likelihood-based tools fit alignments to nucleotide substitution models with three classes of *dN/*dS (synonymous to non-synonymous mutation rate) ratios, usually referred to simply as ω, and compare the fit of competing models to test evolutionary hypotheses. We examined the distributions of the ω*_1_*, ω*_2_*, and ω*_3_* parameters and their associated weights across all orthogroups estimated for the orb-weaving and non-orb-weaving branches of all 4,756 genes tested in this analysis (**Fig. 2a**). We observe a slightly wider overall distribution of ω values across all genes among the non-orb-weaving branches than among the orb-weavers, indicating that a greater frequency of both purifying and diversifying selective events ßhave occurred among the non-orb-weaving species compared to the orb-weavers.

**Figure 2:**
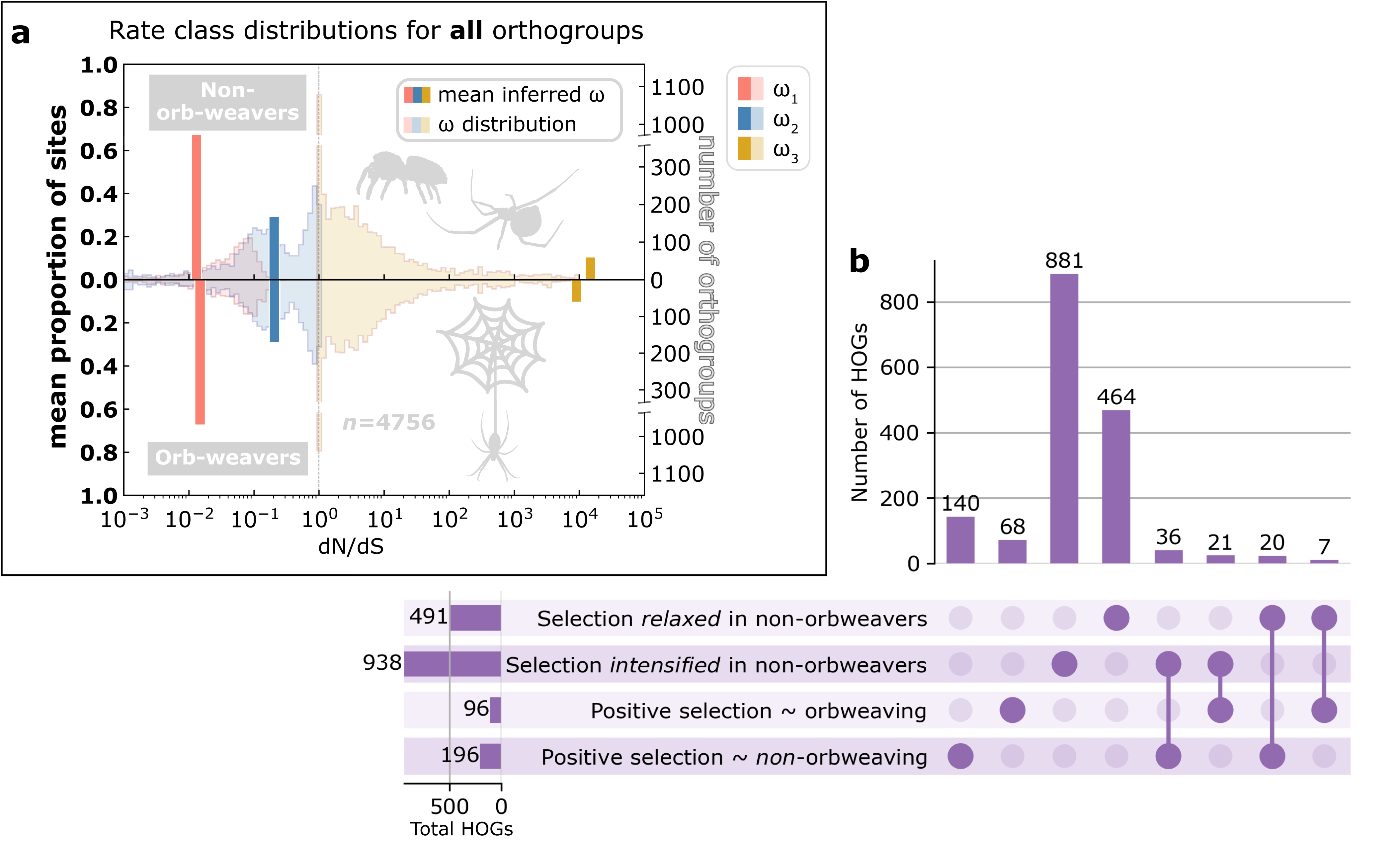
Of 4,756 hierarchical orthogroups (HOGs) tested in four HyPhy analyses, 1,637 showed evidence of selection corresponding to orb-weaving. **(a)** Summary chart showing distributions of inferred values for the three rate classes ω_1_ ≤ ω_2_ ≤ 1 ≤ ω_3_ across all orthogroups (right y-axis) and their corresponding means and proportions (left y-axis) for all 4,756 hierarchical orthogroups (see Methods). Distributions for orb-weavers (bottom) and non-orb-weavers (top) are displayed separately. Values for ω*_1_* (red), ω*_2_* (blue), and ω*_3_* (yellow) for each gene are those estimated by RELAX when fitting to the BS-REL codon substitution model used in both RELAX and BUSTED algorithms. Bars show mean values for ω*_1_*, ω*_2_*, and ω*_3_* across all orthogroups tested, with their heights corresponding to the mean proportion of sites assigned to each rate class across all orthogroups. **(b)** UpSet plot indicating overlapping significant HOGs among the four categories of HyPhy results from RELAX (top two rows) and BUSTED-PH (bottom two rows). The first four bars show counts of genes exclusive to those categories. Silhouette images of spiders are from PhyloPic (phylopic.org).

We found evidence of positive, relaxed, or intensified selection that corresponds to the orb-weaving phenotype (or the non-orb-weaving phenotype) in 1,637 orthogroups out of the 4,756 tested. Among these putative selected genes, of particular interest are those that appeared as significant in more than one of these analyses, which might indicate an even stronger potential relevance to the orb-weaving behavioral program (**Fig. 2b**).

### Relaxation and intensification of selection among non-orb-weaving spiders

RELAX compares evolutionary rates between a test group and a background group for evidence that rates of selection in the test group have relaxed towards neutrality or alternatively that they have intensified towards stronger positive or negative selection, relative to the background branches (Wertheim et al. 2015). Genes that show evidence of relaxed selection are characterized by distributions of ω values closer to 1 in the foreground partition of the phylogeny as compared to the background (**Fig. 3a**). In accordance with the hypothesis that orb-weaving was the ancestral state of all the entelegyne spiders but was lost in the extant non-orb-weaving groups, we specified non-orb-weavers as the foreground partition and searched for genes with evidence of relaxation of selection compared to orthologs in orb-weaving spiders. If the ancestral-with-loss hypothesis of the origin of orb-weaving is correct, we predicted that genes involved in the orb-weaving behavior would show this pattern of selection. We found 491 genes that show evidence of relaxed selection in non-orb-weavers, many of which could be implicated in the orb-weaving behavioral program, such as a novel acetylcholine receptor chaperone (**Fig. 3b**, **Supplementary Table S3**). Acetylcholine is an important neurotransmitter, so the proteins involved in ensuring the proper folding and function of acetylcholine receptors could regulate the coordination of sensory inputs and motor outputs required for orb-weaving. The set of genes that exhibited evidence of relaxation of selection in the non-orb-weavers relative to the orb-weavers showed an enrichment of Gene Ontology (GO) categories (Aleksander et al. 2023) related to neurogenesis and neuronal signal transduction, along with categories known also to be enriched in silk-gland specific gene sets (Chaw et al. 2021), such as transmembrane transporter activity and membrane-related genes (**Fig. 3c**). Construction of an orb web requires a complex complement of silk proteins and silk-producing structures, some of which might not have been as selectively maintained (or, alternatively, as diversified) in spiders that do not weave orb webs.

**Figure 3:**
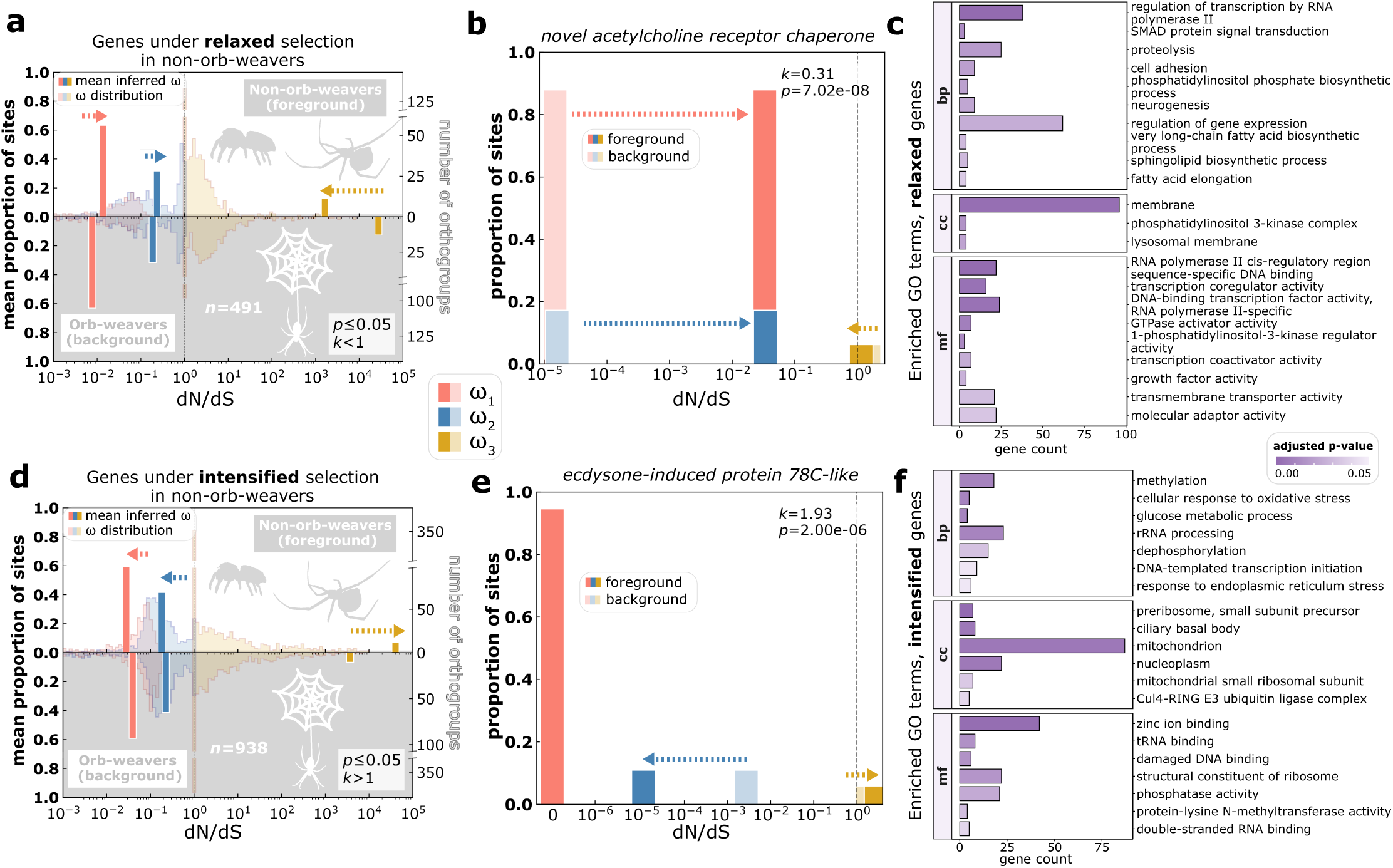
RELAX recovered 491 genes under significantly relaxed selection and 938 genes under significantly intensified selection in non-orb-weavers relative to orb-weavers. Panels (a), (b), (d) and (e) share a central color key. Panels (c) and (f) share a p-value color bar. **(a)** Full ω distributions, mean ω values, and mean proportions of sites assigned to each rate class in each partition for all orthogroups in which selection is significantly relaxed in non-orb-weavers. See Figure 2 legend for a more detailed explanation of this chart style. **(b)** RELAX rate classes and proportions for a gene annotated as *novel acetylcholine receptor chaperone* that shows evidence of relaxed selection in non-orb-weavers. **(c)** Gene Ontology (GO) terms enriched among genes under relaxed selection in non-orb-weavers (Fisher’s Exact Test, *p* ≤ 0.05). GO aspect abbreviations: molecular function (mf), cellular component (cc), biological process (bp). **(d)** Distributions of rate classes for all orthogroups in which selection is significantly intensified in non-orb-weavers. **(e)** RELAX rate classes and proportions for a gene annotated as *ecdysone-induced protein 78C-like* that shows evidence of intensified selection in non-orb-weavers. **(f)** GO terms enriched among genes under intensified selection in non-orb-weavers (Fisher’s Exact Test, *p* ≤ 0.05). Silhouette images of spiders are from PhyloPic (phylopic.org/).

We found an even greater number of orthogroups (938) with evidence of intensified selection among the non-orb-weaving clades compared to their orb-weaving relatives (**Supplementary Table S3**). Genes that experienced an intensification of selection show ω distributions spread further away from the neutral ω = 1 baseline, indicating a greater intensity of positive and/or negative selection in the non-orb-weaving test group compared to the orb-weaving background (**Fig. 3d**). Functional annotation of genes showing this pattern of selection included various promising orb-weaving-related candidates, such as an ecdysone-induced nuclear receptor protein (**Fig. 3e**). Ecdysone is a hormone which plays a role in molting in arthropods, a process which in orb-weaving spiders is generally accompanied by the construction of an orb-shaped “molt web” (Eberhard 2021). Another HOG annotated as *ecdysone receptor-like* also shows an intensification of selection in non-orb-weavers. Furthermore, we found dozens of genes involved in various metabolic processes and cellular responses to oxidative stress with evidence of intensified selection in non-orb-weavers (**Fig. 3f**). These and other categories enriched in this intensified selection set, including mitochondrial genes, could suggest the molecular basis of maintenance of the more active lifestyle practiced by many non-orb-weaving entelegyne spiders, such as the cursorial salticids of the RTA clade that have well developed tracheae for energy-intensive hunting (Schmitz 2005).

### Diversifying selection in entelegyne orb-weavers and non-orb-weavers

Considering instead the hypothesis that orb-weaving evolved multiple times independently, we reasoned that we would find evidence for convergent positive selection in genes related to orb-weaving in clades that today possess the ability to weave an orb web. Thus, we ran the BUSTED-PH test using orb-weavers as the foreground (Kosakovsky Pond 2022). BUSTED-PH is a modification of the original BUSTED algorithm which is optimized to test specifically for evidence of positive selection corresponding to a phenotype of interest (**Fig. 4a**). We found evidence of positive selection associated with the orb-weaving phenotype in 96 genes (**Supplementary Table S4**). For example, a gene showing close homology to Nk-family homeobox protein *ventral nervous system defective* (*vnd*), which in *D. melanogaster* is involved in neurodevelopment (McDonald et al. 1998) and has been shown to be expressed in the developing nervous system in a cob-weaving spider (Schomburg et al. 2020), showed evidence of positive selection only in the orb-weaving partition of its alignment (**Fig. 4b**). Annotation of other identified candidates in this group recovered enrichment for protein phosphorylation, transcription factor TFIID complex, and ATP binding, suggesting that positive selection on genes involved in regulation and phosphorylation signaling cascades could have contributed to the development of complex behavioral programs such as orb-weaving (**Fig. 4c**).

**Figure 4:**
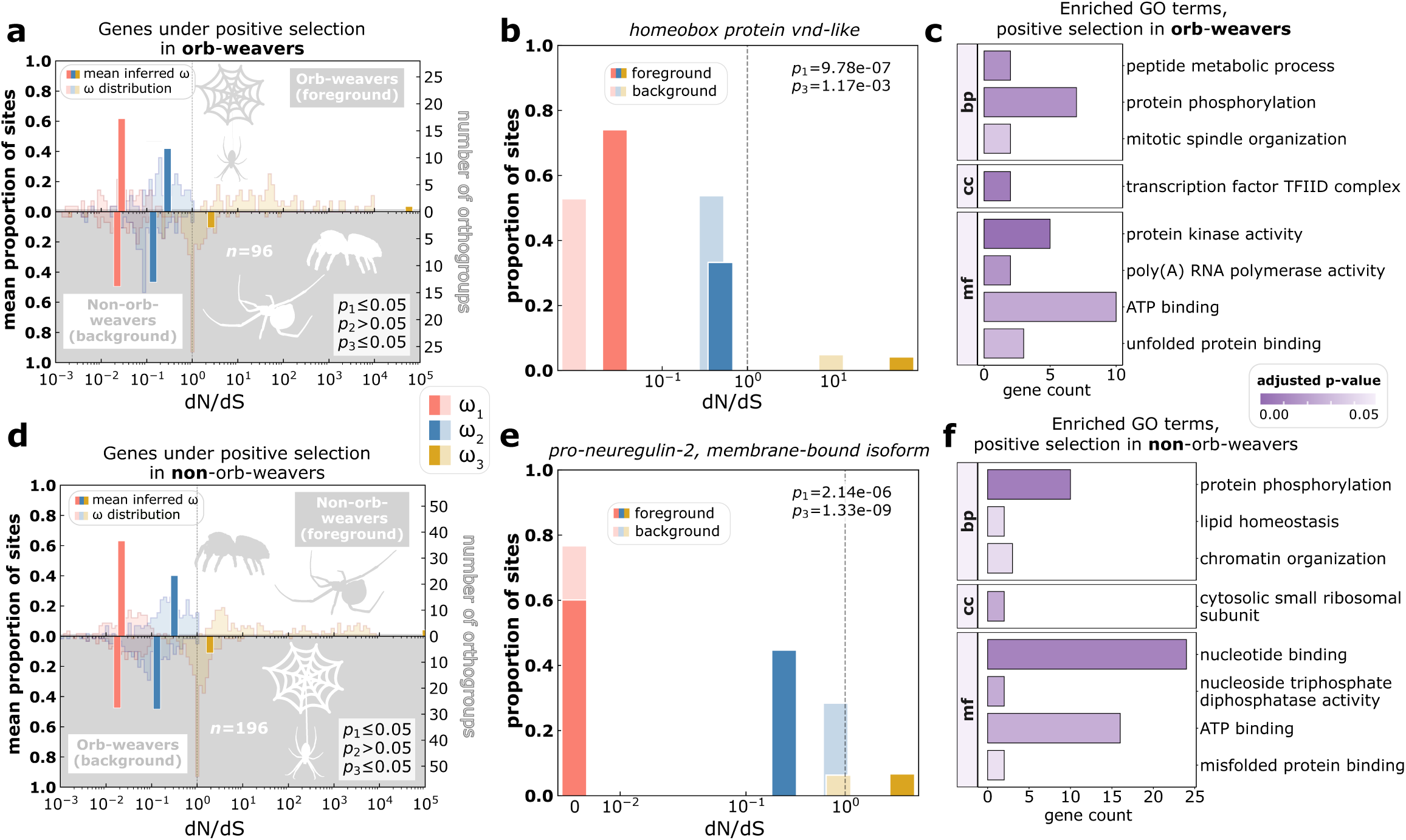
BUSTED-PH recovered 96 genes showing evidence of positive selection associated with the orb-weaving phenotype and 196 genes showing evidence of positive selection associated with the non-orb-weaving phenotype. Panels (a), (b), (d) and (e) share a central color key. Panels (c) and (f) share a p-value color bar. **(a)** Full ω distributions, mean ω values, and mean proportions of sites assigned to each rate class in each partition for all orthogroups that show evidence of positive selection associated with orb-weaving. See Figure 2 legend for a more detailed explanation of this chart style. **(b)** BUSTED-PH rate classes and proportions for a *homeobox protein vnd-like* gene that shows evidence of positive selection only in orb-weavers. **(c)** GO terms enriched among genes positively selected in orb-weavers (Fisher’s Exact Test, p ≤ 0.05). GO aspect abbreviations: molecular function (mf), cellular component (cc), biological process (bp). **(d)** Distributions of rate classes for all orthogroups that show evidence of positive selection associated with non*-*orb-weaving. **(e)** BUSTED-PH rate classes and proportions for a *pro-neuregulin-2 membrane-bound isoform* that shows evidence of positive selection only in non-orb-weavers. **(f)** GO terms enriched among genes positively selected in non*-*orb-weavers (Fisher’s Exact Test, p ≤ 0.05). Silhouette images of spiders are from PhyloPic (phylopic.org/).

The unexpected abundance of genes identified as showing intensified selection in non-orb-weaving lineages (**Fig. 3d–f**) suggested a third hypothesis: that some of the various non-orb web phenotypes and other behavioral traits displayed by the non-orb-weaving entelegyne spiders could have been caused or enabled by changes in ancestral orb-weaving genes (*i.e*., that non-orb-weaving convergently evolved). To test this hypothesis, we ran BUSTED-PH once more on the same dataset, this time selecting the non-orb-weaving lineages as the foreground, to find genes that might have been subject to positive, diversifying selection only in non-orb-weavers (**Fig. 4d**). We discovered 196 such genes (**Supplementary Table S4**), including neural development genes (**Fig. 4e**), genes involved in various metabolic processes, and regulators of gene expression and activity via chromatin organization and post-translational modification (**Fig. 4f**). Here, too, we found strong selection for protein signaling, albeit in a different set of genes.

### Comparing gene loss and gain in orb-weavers versus non-orb-weavers

Although the selection analyses described above provide insight into the evolutionary histories of orthologous genes that might play a role in orb-web construction, they are unable to capture the impact of gene loss and gain on phenotypic innovations across the spider phylogeny. To investigate the association between gene loss or duplication and orb-weaving, we developed a statistical approach to compare the odds of gene loss/duplication between two groups in a phylogeny. For comparative analysis of gene loss, we calculated the odds that a given orb-weaving species lacks the gene and the odds that a non-orb-weaving species lacks the gene. For instance, if 33 of 44 orb-weavers are not represented in a given orthogroup/gene, the odds of loss given the orb phenotype = 3 for that orthogroup. We then calculated the log odds ratio of gene loss in orb-weavers versus non-orb-weavers for each gene (**Fig. 5a**). In the same manner, we calculated the log odds ratio of gene duplication in orb-weavers versus non-orb-weavers for each gene (**Fig. 5b**). Genes with extremely high log odds ratios are significantly more likely to have been lost or duplicated in orb-weavers, and those genes with low log odds ratios would be more likely lost or duplicated in non-orb-weavers.

**Figure 5:**
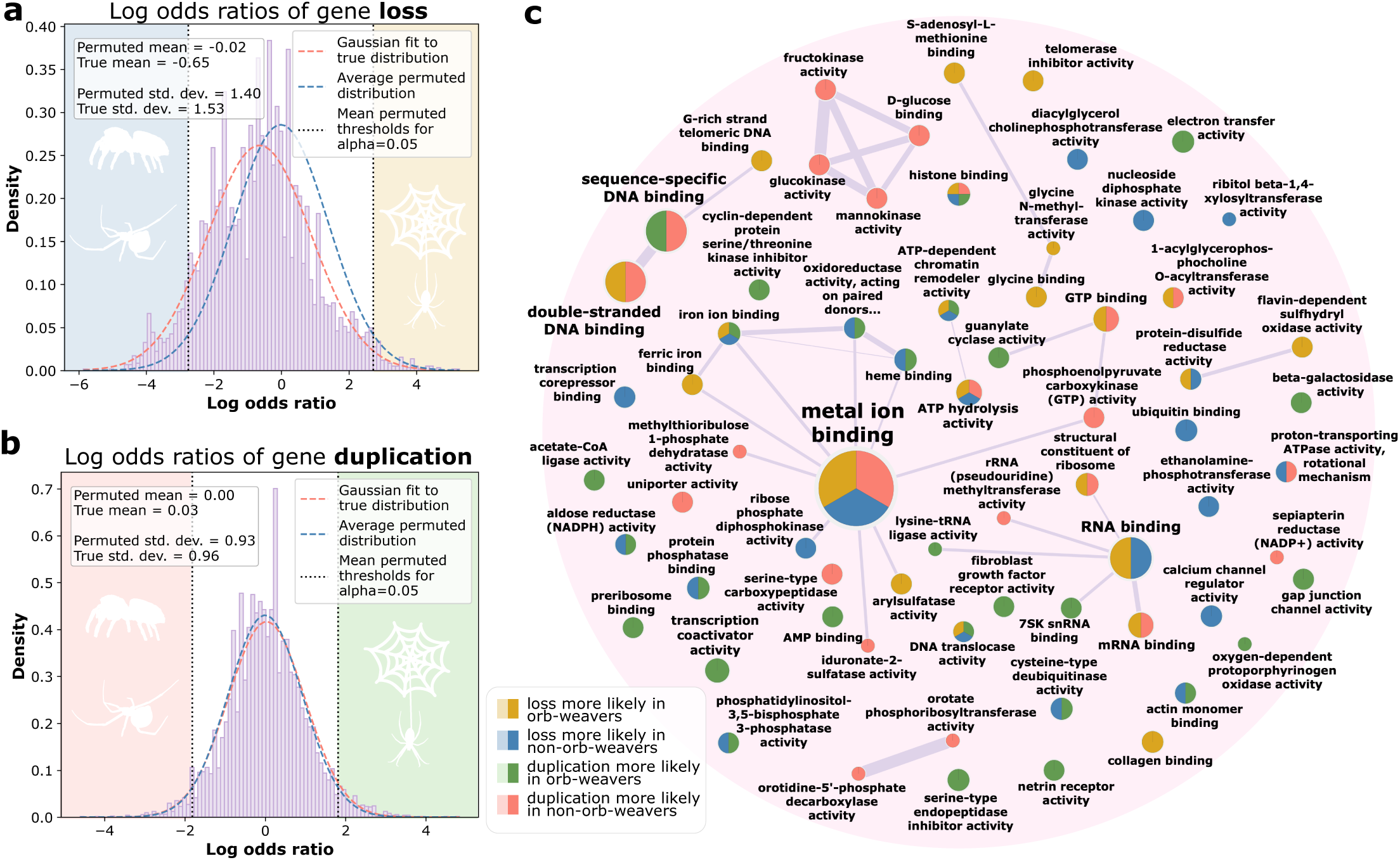
Permulation-based analysis of differences in odds of gene loss and duplication between orb-weavers and non-orb-weavers. All panels share the color key at bottom center. **(a)** Distribution of log odds ratios (LORs) of loss in each orthogroup among orb-weavers vs. non-orb-weavers. The average 95% confidence interval of all 10,000 permulated distributions was used as the threshold for significance (black dotted lines). Yellow region, LOR > 2.718, loss more likely in orb-weavers; blue region, LOR < –2.758, loss more likely in non-orb-weavers. **(b)** Distribution of log odds ratios of duplication in each orthogroup among orb-weavers vs. non-orb-weavers. Green region, LOR > 1.812, duplication more likely in orb-weavers; red region, LOR < –1.819, duplication more likely in non-orb-weavers. Silhouette images of spiders are from PhyloPic (phylopic.org/). **(c)** Network plot showing significantly enriched categories of molecular functions. Sizes of category nodes and labels correspond to size of the gene set associated with that GO term. Widths of edges correspond to the similarity coefficient (Jaccard index) between connected gene sets/nodes. Nodes show equal wedges indicating in which of the four categories that GO term was enriched; sizes do not reflect relative contributions.

We then used a permutation-based approach to test two hypotheses: (1) that non-orb-weavers in general are more likely to have experienced gene loss relative to orb-weavers, in line with the ancestral-with-loss model, and (2) that orb-weavers are more likely to have duplicated genes than non-orb-weavers, as might be predicted if the trait evolved convergently. Under null hypotheses that the odds of gene loss and gain are not significantly different between orb-weavers and non-orb-weavers across all genes, the true distribution of log odds ratios should not be significantly different from distributions in which the foreground and background groups were assigned randomly, with attention to phylogenetic relatedness. To generate null distributions of randomized phenotype assignments while preserving potentially relevant phylogenetic information, we used a “permulations” approach (Saputra et al. 2021). This method uses a combination of phylogenetic simulations and permutations to generate null phenotypes that account for the correlation structure in the data introduced by the relatedness of the species. We generated 10,000 permulated phenotype vectors and used these to calculate the parameters of 10,000 null log odds ratio (LOR) distributions, counting the number of times the mean of the null distribution exceeded the mean of the true distribution (for gene duplication) or was less than that of the true distribution (for gene loss).

We found no evidence that orb-weaving spiders are in general more likely to have multiple copies of shared genes than their non-orb-weaving relatives (*p* = 0.3254) nor that non-orb-weavers are more likely overall to have lost genes (*p* = 0.1854). The permulated null distributions provided a reference from which we determined thresholds of log odds ratios more extreme than would be expected under a null hypothesis that the orb-weaving phenotype has no bearing on the odds of a particular gene having been lost or duplicated. Using the accumulated parameters from the 10,000 permulated LOR distributions, we calculated an average 95% confidence interval, representing the expected range of log odds ratios of loss and gain under the null hypotheses. The permulated confidence interval was (–2.758, 2.718) for the log odds ratio distribution of gene loss (**Fig. 5a**) and (–1.819, 1.812) for the log odds ratio distribution of gene duplication (**Fig. 5b**). For genes with log odds ratios that fall outside of this range, the odds of loss or duplication are significantly different in orb-weavers compared to non-orb-weavers (**Table Supplementary S5**).

These gene sets showed enrichment for various neural and silk-related GO terms (**Fig. 5c, Supplementary Fig. S1**) and showed considerable overlap with the significant results from the selection analyses and genes that might be silk-gland specific (**Fig. 6**, **Supplementary Table S6**). Contrary to our expectations, none of the genes identified as having a history of selective pressure or copy number change corresponding to orb-weaving were identifiable as spider silk genes (spidroins). Because we did not investigate the ancestral state of the genes tested in this analysis, it should be mentioned that genes designated as “lost” in one group could conceivably be designated as “novel” genes in the other group, which never existed in their common ancestor, and thus were never actually lost by the former. Similarly, genes we have described as more likely to be “duplicated” in one group could in fact have been present in multiple copies in the ancestor, but some of those ancestral copies were in fact lost in the other group. Despite these uncertainties, we nevertheless consider these genes with significant differences in copy number between orb-weaving and non-orb-weaving spiders to be worthy of further inquiry into their behavioral relevance.

**Figure 6:**
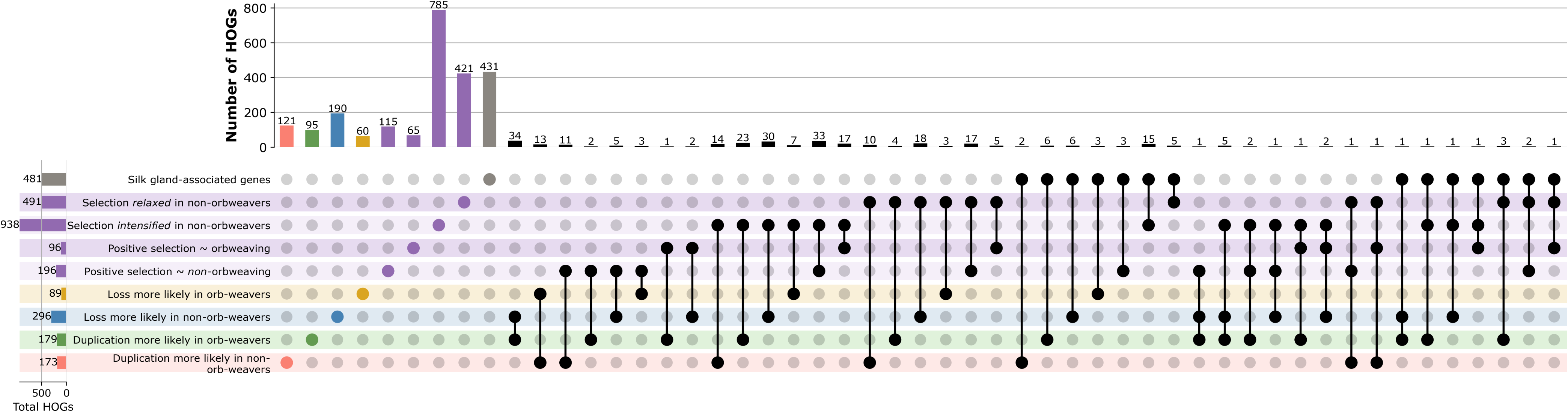
UpSet plot indicating counts of HOGs found as significant in multiple analyses performed in this study and of HOGs with a high degree of homology to known silk-gland-associated genes in *Argiope argentata* (Chaw et al. 2021). The first nine bars show counts of genes exclusive to each category. The remaining bars correspond to genes that are significant in multiple analyses and/or show homology to silk gland genes. See Supplementary Table S2 for a full list of these orthogroups.

## Discussion

Here we used existing and original statistical approaches to probe the molecular foundations and evolutionary origins of complex behaviors at the gene level. Where previous maximum likelihood studies debated the evolutionary history of orb-weaving based solely on unstable phylogenetic relationships among species, we interrogated that history directly by searching for the signatures it has left on specific genes and loci which give rise to the behavior. By interrogating these individual genetic elements showing evidence of association with orb-weaving, we obtain a more detailed picture of the evolutionary forces that underlie the emergence of such a precisely organized, hierarchical behavior. Our findings suggest that different genes involved in modern-day orb-weaving could have evolved under different, even contradictory selection regimes. Genes that give rise to some aspects of the behavior might be ancestral; others might have arisen convergently; and still more appear to have been a starting point for further innovation and elaboration, leading to the myriad web types and lifestyles we observe among entelegyne spiders today.

As a product of this analysis, we generated a dataset of genes which could be relevant to the orb-weaving phenotype, of which we highlight a few that we consider especially promising for future experimental studies (**Table 1**). Genes with potential relevance to neural development and function, such as a *semaphorin-2A-like* gene that is both relaxed in non-orb-weavers and more likely to be lost in non-orb-weavers, are obvious choices for genetic intervention in behavioral assays. Analysis of gene copy number in particular generated several genes involved in G-protein signaling, including receptors and downstream proteins involved in inositol phosphate signaling. Both orb-weavers and non-orb-weavers were enriched for different proteins in these signaling pathways. Neuromodulation strongly shapes behavior and predominantly operates through G-protein signaling (Muir et al.). Gene duplication in these systems provides an efficient pathway towards evolving altered signaling and, subsequently, behavior.

**Table 1:** Highlighted orthogroups showing evidence of selection for orb-weaving.*

| Gene description | Orthogroup ID | Occ. | RELAX,<br>*non-orb-weavers* FG |  |  | BUSTED-PH, *orb-weavers* FG |  | BUSTED-PH, *non-orb-weavers* FG |  | Odds ratio test<br>Orb:non-orb LOR |  | Test results summary |
| --- | --- | --- | --- | --- | --- | --- | --- | --- | --- | --- | --- | --- |
|  |  |  | p-value | LR | k | p-values | LRs | p-values | LRs | loss | duplication |  |
| novel acetylcholine receptor chaperone | N5.HOG0066983 | 94 | 7.02E-08 | 29.1 | 0.307 | *NS* | *NS* | *NS* | *NS* | *NS* | *NS* | selection relaxed in non-orb-weavers |
| ecdysone-induced protein 78C-like | N5.HOG0048857 | 97 | 0.00570 | 7.64 | 1.25 | *NS* | *NS* | *NS* | *NS* | *NS* | *NS* | selection intensified in non-orb-weavers |
| homeobox protein *vnd*-like | N5.HOG0069577 | 81 | *NS* | *NS* | *NS* | 9.78E-07;<br>0.5; 1.17E-03 | 26.3; -<br>10.3; 20.2 | *NS* | *NS* | *NS* | *NS* | positive selection in orb-weavers |
| pro-neuregulin-2, membrane-bound isoform | N5.HOG0028049 | 93 | *NS* | *NS* | *NS* | *NS* | *NS* | 2.14E-06;<br>0.5; 1.33E-09 | 24.7;<br>0; 50.1 | *NS* | *NS* | positive selection in non-orb-weavers |
| semaphorin-2A-like | N5.HOG0025053 | 87 | 1.24E-03 | 10.4 | 0.898 | *NS* | *NS* | *NS* | *NS* | -3.11 | *NS* | selection relaxed in non-orb-weavers;<br>more likely lost in non-orb-weavers |
| homeobox protein abdominal-B-like | N5.HOG0044354 | 75 | *NS* | *NS* | *NS* | *NS* | *NS* | *NS* | *NS* | *NS* | 1.89 | more likely duplicated in orb-weavers |
| male-specific lethal 1 | N5.HOG0054505 | 84 | 2.19E-05 | 18.0 | 0.846 | *NS* | *NS* | *NS* | *NS* | 3.89 | *NS* | selection relaxed in non-orb-weavers;<br>more likely lost in orb-weavers |
| male-specific lethal 3 | N5.HOG0053389 | 75 | *NS* | *NS* | *NS* | *NS* | *NS* | 0.0281;<br>0.5; 1.34E-03 | 5.76; 0;<br>19.8 | -4.14 | *NS* | positive selection in non-orb-weavers;<br>more likely lost in non-orb-weavers |
| males absent on the first | N5.HOG0045324 | 84 | 7.78E-05 | 15.6 | 0.890 | *NS* | *NS* | *NS* | *NS* | *NS* | *NS* | selection relaxed in non-orb-weavers |
| inositol phosphate kinase 2 | N5.HOG0060564 | 93 | 2.68E-04 | 13.3 | 0.587 | *NS* | *NS* | *NS* | *NS* | *NS* | *NS* | selection relaxed in non-orb-weavers |
| sphingosine-1-phosphate phosphatase 2-like | N5.HOG0067221 | 89 | 5.35E-04 | 12.0 | 0.425 | *NS* | *NS* | *NS* | *NS* | *NS* | *NS* | selection relaxed in non-orb-weavers |
| inositol-tetrakisphosphate 1-kinase-like | N5.HOG0060995 | 97 | *NS* | *NS* | *NS* | 4.88E-04;<br>0.440;<br>3.10E-02 | 13.9;<br>0.257;<br>12.3 | *NS* | *NS* | *NS* | *NS* | positive selection in orb-weavers |
| neurogenic locus notch homolog protein 1-like | N5.HOG0014651 | 75 | 6.40E-04 | 11.7 | 0.797 | *NS* | *NS* | *NS* | *NS* | *NS* | *NS* | selection relaxed in non-orb-weavers |
| spondin-1-like | N5.HOG0027314 | 94 | 1.55E-05 | 18.7 | 0.347 | *NS* | *NS* | 0.0472;<br>0.444;<br>8.204E-5 | 4.72;<br>0.239;<br>26.2 | *NS* | *NS* | selection relaxed in non-orb-weavers; positive<br>selection in non-orb-weavers; enriched in silk<br>glands |
| phosphatidylinositol 3-kinase 92E | N5.HOG0050702 | 95 | 4.76E-08 | 29.8 | 0.916 | *NS* | *NS* | *NS* | *NS* | *NS* | *NS* | selection relaxed in non-orb-weavers |
| netrin receptor UNC5C | N5.HOG0028124 | 55 | *NS* | *NS* | *NS* | *NS* | *NS* | *NS* | *NS* | *NS* | 2.56 | more likely duplicated in orb-weavers |
| progranulin-like | N5.HOG0033903 | 66 | *NS* | *NS* | *NS* | *NS* | *NS* | *NS* | *NS* | -3.04 | 2. | more likely lost in non-orb-weavers;<br>more likely duplicated in orb-weavers |
| Schwannomin interacting protein 1 | N5.HOG0037962 | 95 | NS | NS | NS | NS | NS | NS | NS | NS | 2.33 | more likely duplicated in orb-weavers |
| innexin inx1-like | N5.HOG0055111 | 56 | NS | NS | NS | NS | NS | NS | NS | NS | 2.26 | more likely duplicated in orb-weavers |
| CB1 cannabinoid receptor-interacting protein 1-like | N5.HOG0037768 | 83 | 7.89E-05 | 15.6 | 0.545 | NS | NS | NS | NS | NS | NS | selection relaxed in non-orb-weavers |
| G-protein coupled receptor moody-like | N5.HOG0042919 | 94 | 9.75E-04 | 10.9 | 0.723 | NS | NS | NS | NS | NS | NS | selection relaxed in non-orb-weavers |
| phosphatidylinositol 4-kinase type 2-beta-like | N5.HOG0038628 | 98 | 4.25E-06 | 21.2 | 0.700 | NS | NS | NS | NS | NS | NS | selection relaxed in non-orb-weavers |
| inositol 1,4,5-triphosphate receptor associated 1-like | N5.HOG0030773 | 97 | 3.06E-03 | 8.77 | 0.885 | NS | NS | NS | NS | NS | NS | selection relaxed in non-orb-weavers |
| G protein pathway suppressor 2-like | N5.HOG0059007 | 96 | 2.26E-02 | 5.20 | 0.916 | NS | NS | NS | NS | NS | NS | selection relaxed in non-orb-weavers |
| synaptobrevin homolog YKT6-like | N5.HOG0042219 | 98 | 2.41E-02 | 5.09 | 0.882 | NS | NS | NS | NS | NS | NS | selection relaxed in non-orb-weavers |
| GTP-binding protein REM 1-like | N5.HOG0022784 | 76 | 1.24E-07 | 28.0 | 0.295 | NS | NS | NS | NS | NS | NS | selection relaxed in non-orb-weavers |
| macoillin-like | N5.HOG0059145 | 98 | 3.87E-02 | 4.27 | 0.904 | NS | NS | NS | NS | NS | NS | selection relaxed in non-orb-weavers |
| transcription factor *cwo* | N5.HOG0058528 | 95 | 2.65E-03 | 9.03 | 0.457 | NS | NS | NS | NS | NS | NS | selection relaxed in non-orb-weavers |
| inositol monophosphatase 1-like | N5.HOG0056026 | 98 | NS | NS | NS | NS | NS | 8.16E-10;<br>0.5;<br>5.97E-04 | 40.5;<br>0;<br>21.7 | NS | NS | positive selection in non-orb-weavers |
| syntaxin-binding protein *Rop* | N5.HOG0041449 | 97 | NS | NS | NS | NS | NS | 8.50E-05;<br>0.5;<br>6.39E-05 | 17.4;<br>0;<br>26.7 | NS | NS | positive selection in non-orb-weavers |
| tyrosine-protein phosphatase non-receptor type 5-like | N5.HOG0062144 | 90 | NS | NS | NS | 2.04E-03;<br>0.299;<br>1.28E-04 | 11.0;<br>1.03;<br>25.2 | NS | NS | NS | NS | positive selection in orb-weavers |
| mesencephalic astrocyte-derived neurotrophic factor homolog | N5.HOG0055615 | 98 | 3.76E-02 | 4.32 | 0.919 | NS | NS | NS | NS | NS | -2.72 | selection relaxed in non-orb-weavers; more likely duplicated in non-orb-weavers |

\* **Gene descriptions** are from the annotated *U. diversus* ortholog of the gene (Miller et al. 2023). **Occ.** refers to the *occupancy* of the orthogroup (how many species have at least one gene in the orthogroup). For the BUSTED-PH tests, the three p-values and three likelihood ratios correspond to the three hypothesis tests: (1) for positive selection in the foreground; (2) for positive selection in the background; and (3) for difference between the foreground and background distributions. For a significant result, $p_1 \leq 0.05$ , $p_2 > 0.05$ , and $p_3 \leq 0.05$ . For the odds ratio tests, a negative log odds ratio (LOR) indicates that loss/duplication is more likely in non-orb-weavers, whereas a positive LOR means loss/duplication is more likely in orb-weavers. Numerical values of statistics/parameters are only shown for statistically significant results. Individual genes belonging to each spider species in the orthogroup and complete test results can be found in **Supplementary Tables S2-5**. Additional significant genes of particular interest from the odds ratio test analyses are compiled in **Supplementary Table S7**.

Several putative homeotic genes, which are known for their role in body plan development, appear among the significant results, particularly in the set of genes more likely to be lost in non-orb-weaving spiders. Among these, any genes that could conceivably be involved in developing anatomical features relevant to orb-weaving would be excellent candidates as well. For example, an orthogroup annotated as the Hox gene *Homeobox protein abdominal-B-like* (*abd-B*) showed significance in the test for differential odds of duplication. Because spiders, like mammals, have two Hox clusters owing to an ancestral whole genome duplication (Aase-Remedios et al. 2023), this result could suggest either that orb-weavers are more likely to have *retained* both ancestral copies of abdominal-B, or that they are more likely to have further duplicated the gene to provide fodder for additional adaptation. *Abd-B* also shows homology to a gene known to be enriched in spider silk glands.

We take a particular interest in genes identified by these analyses that might have some role in sexual development because in many orb-weaving species, the orb-weaving behavior is itself sexually dimorphic (adult males do not typically weave orb webs). The evolution of orb-weaving may thus have been intertwined with the evolution of spider sexual dimorphism in general. Spider genes orthologous to the *Drosophila* male-specific lethal 1, male-specific lethal 3, and males absent on the first (MSL1, MSL3, MOF), which together form substantial part of the MSL complex required for dosage compensation in male flies (Babosha et al. 2023), all returned significant results in at least one of the HyPhy analyses, and each MSL gene was more likely to have been lost in one of the two phenotypic groups. Though the copy number results could be skewed by imbalanced representation of the sexes among the specimens from which these transcriptomes were derived, the signatures of selection we found are likely valid.

We also provide a list of orthogroups which appeared as significant in multiple analyses, which include several nervous system and neuron development-related genes, genes associated with G-protein coupled receptor activity, and sex-related genes (**Supplementary Table S6**). We also include genes which we have identified as having close homology with known silk-gland-specific transcripts in another orb-weaver, *Argiope argentata* (Chaw et al. 2021). Many of the genes appearing as significant in three or more of these tests are as yet unannotated and may be responsible for spider-specific functions which have yet to be discovered. If any of the genes identified by these analyses are validated experimentally as having an essential role in the orb-weaving behavior, it could lend convincing support to one of the competing hypotheses about the evolutionary origins of the behavior.

In this study, we implemented a new approach to exploring the origins of orb-weaving by testing evolutionary hypotheses on several thousand protein-coding genes. Given the complex, time-dependent and developmental stage-specific nature of the orb-weaving behavioral program, it is likely that much of the relevant evolutionary change that has enabled orb-weaving and/or its loss has occurred in regulatory elements and other non-coding genetic elements. Because of the nature of the HyPhy selection tests, which are based on non-synonymous and synonymous mutation rates, and the requirements of OrthoFinder, which expects protein sequences, these more elusive regions of the genome were not included in this analysis and constitute an interesting line of inquiry for future studies. Beyond the promising orb-weaving-related gene candidates identified here, our approach and extensive dataset of orthologous spider genes provide a framework for examining the evolutionary histories and genetic underpinnings of other spider behaviors and complex behaviors in general.

## Methods

### Transcriptome data collection and processing

We acquired 573 publicly available spider transcriptomes via the Nucleotide, SRA, BioProject and BioSample databases from the National Center for Biotechnology Information (Sayers et al. 2024), with a sizable number coming from the 1,000 Silkomes Project (Arakawa et al. 2022) (**Supplementary Table S1**). We chose transcriptomes in place of genomes because of the relative dearth of annotated spider genomes at the commencement of this study, though that number has since increased meaningfully. We clustered raw transcript sequences to remove redundancy with CD-HIT (Li and Godzik 2006), then identified open reading frames at least 100 amino acids long using TransDecoder.LongOrfs (Haas 2022). We submitted these ORFs as a BLAST-P (Camacho et al. 2009) query against the UniProt database (The UniProt Consortium 2025) to identify any strong homology to known proteins (e-value < 1E-5). We used TransDecoder.Predict to predict coding sequences from the long ORFs, with the option to retain any ORF with a BLAST-P match in the final output. We analyzed the resulting peptide sequences for redundancy and completeness by searching for the presence and uniqueness of the arachnid-lineage set of Benchmarking Universal Single Copy Ortholog (BUSCO) genes, version odb10 (Manni et al. 2021). We filtered the 573 transcriptomes we had initially curated for quality using a threshold of no less than 90% total complete BUSCOs and no more than 30% duplicated BUSCOs, with 3 exceptions: *Antrodiaetus roretzi*, which had 86% total complete BUSCOs but we included as it was the highest-quality representative of *Mygalomorphae*, an important ancestral family; and two species of the family *Deinopidae*, which had 88-89% complete BUSCOs but we determined to be essential to the analysis as cribellate “orb-weavers” who are sister to the model cribellate orb-weaver *U. diversus*. The deinopids’ status as orb-weavers is not universally affirmed; also known as net casters, they weave an orb-like web, which they suspend between their front legs to physically cast upon their prey (Coddington 1986). For the purpose of this study, we considered them to be cribellate orb-weavers (see *Orb-weaving phenotype designation* below for more details). This filtering left a total of 102 spider species. We downloaded protein sequences for *Drosophila melanogaster*, which we chose as a non-spider outgroup for the orthology search, from FlyBase (dmel-all-translation release 6.51) (Öztürk-Çolak et al. 2024).

### Orthology search and pre-testing pipeline

We fed the 103 species’ protein sequences into OrthoFinder v2.5.5 (Emms and Kelly 2019) with default settings. After initially generating orthogroups, orthologs, and a resolved species tree, we analyzed and manually edited the species tree using Interactive Tree of Life (Letunic and Bork 2024) to better accord with the most recent, best resolved and supported spider tree of life at the time (Kulkarni et al. 2023) (**Fig. 1a-b**). We moved five leaves of the OrthoFinder-generated tree to achieve this. We then re-ran the final steps of the OrthoFinder pipeline using the “-ft” option with the edited species. For the tests of positive and relaxed selection, we filtered the list of phylogenetic hierarchical orthogroups (HOGs) from the fifth node (N5), which included all entelegyne spiders and excluded *Drosophila* and four non-entelegyne spiders, to those HOGs with an occupancy of at least 75 species and in which the spider *U. diversus*, the species whose annotations we used for ontology analysis, was represented by at least one gene. We then used the filtered list of 4,756 entelegyne HOGs to generate individual fasta files for each HOG containing the coding sequences of all genes in the HOG via a custom script.

To prepare the orthogroup fasta files for evolutionary hypothesis testing, we first subjected the N5.HOG files to pre-alignment quality filtering using PREQUAL v1.02 (Whelan et al. 2018) and then generated multiple sequence alignments for each gene using the alignSequences program of MACSE v2.07 with default settings (Ranwez et al. 2018). We inferred a gene tree for each orthogroup using IQ-TREE v2.3.6 (Minh et al. 2020). We performed an initial run of the HyPhy (v2.5.70) batch language-based BUSTED (v4.6) test for episodic diversifying selection on the alignments and corresponding tree files with the --multiple-hits Double+Triple and --error-sink Yes” settings, but with no foreground specified, to generate the error-filtered alignment BUSTED produces when using the --error-sink flag (Murrell et al. 2015; Kosakovsky Pond et al. 2020). We parsed these error-filtered alignments and corresponding edited trees using HyPhy error-filter and labeled any branches on the trees corresponding to genes from orb-weaving species as foreground using the HyPhy LabelTrees tool.

### Orb-weaving phenotype designation

We classified all araneids as orb-weavers aside from the three-dimensional dome-web weaver *Cyrtophora cicatrosa* (Rao and Poyyamoli), the bolas spiders *Ordgarius hobsoni* and *Ordgarius sexspinosus* (Tanikawai 1997), and *Cyrtarachne bufo*, which belongs to a genus sister to the bolas spider that builds a reduced orb web with transitional bolas-like functionality (Cartan and Miyashita 2000). We classified all tetragnathids as orb-weavers. We included the cribellate orb-weaver *U. diversus* and two species of ogre-faced spiders of the sister clade *Deinopidae*, whose partial orb-like web which it wields as a net is thought to be synapomorphic to the uloborid cribellate orb (Coddington 1986; Garrison et al. 2016), as the final three orb-weaving species. We designated all remaining spider species as non-orb-weavers.

### Testing for positive and relaxed selection

To test for evidence of relaxation of selection in the 4,756 HOGs, we used RELAX (Wertheim et al. 2015). RELAX uses the basic unrestricted codon model as its null model and compares it to an alternative in which the rate classes are modified by an additional parameter *k.* If the LRT is significant, a value of *k* < 1 indicates the degree of relaxation of selection in the foreground branches relative to the background, whereas *k* > 1 indicates an intensification of selection. We ran RELAX v4.5 on all the alignments and corresponding trees with non-orb-weaving species as the foreground using the following options: --test Unlabeled branches --multiple-hits Double+Triple --models Minimal --srv Yes ENV=“TOLERATE_NUMERICAL_ERRORS=1;”.

We used BUSTED-PH (Kosakovsky Pond 2022), a stand-alone analysis built upon BUSTED (Murrell et al. 2015) to test for evidence of positive selection associated with a particular trait or phenotype, in this case orb-weaving. BUSTED-PH performs three likelihood ratio tests (LRT): (1) for positive selection in the foreground; (2) for positive selection in the background; and (3) for difference between the foreground and background distributions, thus providing insight about the relevance of the phenotype to patterns of selective pressure across the phylogeny. We performed BUSTED-PH v0.2 on the same set of genes and trees, first designating orb-weaver branches as the test group, then designating non-orb-weaver branches as the test group, with default settings and the same environment variable as described for RELAX.

We parsed genes with significant results from the BUSTED-PH and RELAX JSON files, filtering out results with extreme ω values (mean ω > 10,000, calculated as ω_1_ • *P*_1_ + ω_2_ • *P*_2_ + ω_3_ • *P*_3_ where ω_1_ ≤ ω_2_ ≤ 1 ≤ ω_3_ and *P*_<1,2,3>_ are the proportion of sites belonging to each rate class), as extremely large ω values represent unrealistic evolutionary processes and are most likely the result of alignment or sequencing errors (Selberg et al. 2025). We determined the LOC identifiers for the *U. diversus* gene(s) present in each significant orthogroup to use for functional analysis. For RELAX, significantly relaxed genes had *p* ≤ 0.05 and *k* < 1, and significantly intensified genes had *p* ≤ 0.05 and *k* > 1. For BUSTED-PH, significant genes had a foreground p-value ≤ 0.05, a background p-value > 0.05, and a shared p*-*value < 0.05. Plots of rate class distributions and means (**Figs. 2–4**) were visualized using matplotlib (Hunter 2007).

### Gene loss and duplication analysis

To analyze differences in the odds of gene loss between orb-weavers and non-orb-weavers, we used a larger subset of the N5 HOGs from the 98 species of entelegyne spiders (**Supplementary Table S2**) and filtered for minimum occupancy of 50. We chose a lower occupancy threshold here than in the HyPhy-based analyses to serve the goal of identifying genes that are missing in some species, which makes lower occupancy orthogroups particularly relevant. For the test of differential odds of loss only, we also set a maximum occupancy threshold of 95 to ensure that a meaningful difference in odds of loss was possible between the two test groups. For this test, we used the results from OrthoFinder, specifically the GeneCount file, which gives the number of genes per species in each orthogroup, and which can be generated using the OrthoFinder analysis tool orthogroup_gene_count.py from the OrthoFinder GitHub (Emms 2026). We first calculated the odds that a given orb-weaving (foreground) species lacks the gene and the odds that a non-orb-weaving (background) species lacks the gene. We scaled these odds according to the BUSCO scores (**Supplementary Table S1**) for each species to account for the impact of the variation in completeness and redundancy of the transcriptomes. We then determined the log odds ratio (LOR) between the two groups for each orthogroup, using the Haldane-Anscombe correction (Anscombe 1956; Haldane 1956) to account for any odds equaling zero, which would result in undefined LORs:

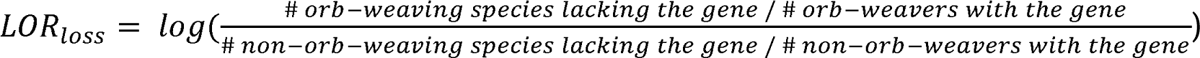

This procedure was repeated to analyze the relative odds of duplication, defined as having the gene in more than one copy:

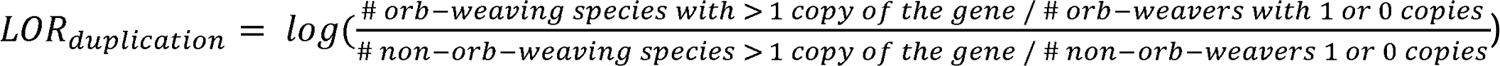

Because we set a maximum occupancy for the loss test but not the duplication test, HOGs in which two or fewer species were missing the gene were excluded from the loss test but included the duplication test. Thus, 7,875 HOGs were evaluated in the duplication test, and 5,269 HOGs were used in the loss test. The true distributions of LORs for loss and duplication across all genes are found in **Figures 5a–b**.

We then generated 10,000 permuted and simulated phylogenies (“permulations”) and associated phenotype vectors using the categoricalPermulations function from the RERconverge package (Saputra et al. 2021). We provided as input the list of orb-weaving species and the resolved species tree with branch lengths generated by OrthoFinder from the original dataset. For each permulated phenotype designation, we re-calculated the LORs for duplication and loss across all genes. We counted how many times the mean of the permulated LOR distribution was smaller than (for the loss test) or exceeded (for the duplication test) the mean of the true distribution. From these counts, we determined the probability that the mean of the true distribution of log odds ratios between orb-weavers and non-orb-weavers was significantly greater (for loss) or less (for duplication) than what would be expected by random chance, given the phylogenetic structure of the data.

For each permulation, we modeled the (null) LOR as a single Gaussian distribution and recorded its mean, standard deviation, and 95% confidence interval thresholds (**Supplementary Fig. S2**). For each of the two tests (loss and duplication), we used the average permulated confidence interval to set benchmark thresholds upon which to judge the extremeness of the true LORs. The orthogroups for which the LORs of loss or duplication in orb-weavers versus non-orb-weavers was outside of the average permulated confidence interval were considered significant. We filtered those orthogroups for the presence of at least one gene from *U. diversus*, both to enable ontology enrichment analysis and to provide some additional assurance that the differences recorded were connected to the orb-weaving phenotype itself, and not just araneoid-specific genes, given that the vast majority of orb-weavers are araneids or tetragnathids. The distributions were visualized using matplotlib and seaborn (Hunter 2007; Waskom 2021).

We noticed that the true LOR distributions appeared potentially multimodal and that the distributions of permulated means were consistently trimodal. We fit the distributions of LORs of loss and duplication to single, double, triple, and quadruple Gaussian models and found the parameters which maximized the likelihood of each model using scipy.optimize.minimize with the “COBYLA” method (Jones et al. 2001). Using the Bayes Information Criterion for each model, we determined that the triple Gaussian was the best fit to both distributions. On this basis, we repeated the permulation test, but this time we fit each null distribution to a triple Gaussian model and counted the null distributions for which each of the three means was smaller than (for loss) or exceeded (for duplication) the corresponding triple Gaussian mean in the true distribution (**Supplementary Fig. S3**). We still did not find any significant differences between the parameters of the true distributions and those of the null distributions when modeled as triple Gaussian (for loss, *p*_μ1_ *=* 0.0933, *p*_μ2_ *=* 0.2893, and *p*_μ3_ *=* 0.1383; for duplication, *p*_μ1_ *=* 0.1169, *p*_μ2_ *=* 0.3424, and *p*_μ3_ *=* 0.6199). Full details of this analysis can be found on GitHub: github.com/GordusLab/orb-selection).

### Ontology enrichment analysis

We used topGO R package v2.58.0 (Alexa et al. 2024) to perform GO enrichment using Fisher’s Exact Test on the *U. diversus* gene IDs from the orthogroups with significant results from each of the above analyses (RELAX; BUSTED-PH, orb-weaver foreground; BUSTED-PH non-orb-weaver foreground; loss odds ratio test; duplication odds ratio test) against the full set of *U. diversus* genes tested in each analysis. We adapted the topGO pipeline from *topGO_pipeline* (Liew 2018) on GitHub. The GO barplots (**Figs. 3c,f; 4c,f**) and topGO_pipeline further used R packages forcats (Wickham 2025), ggpubr (Kassambara 2025), ggplot2 (Wickham 2016), and dplyr (Wickham et al. 2025). We generated the network plot (**Fig. 5c**) using the EnrichmentMap app v3.5.0 (Merico et al. 2010) in Cytoscape (Shannon et al. 2003). We made UpSet plots (Lex et al. 2014) using the python UpSetPlot package (Nothman 2024). We generated the list of *U. diversus* silk gland genes by finding the best tBLASTn hit in the *U. diversus* genome of each of the silk-gland-specific genes identified in *Argiope argentata* by (Chaw et al. 2021).

## Acknowledgements

We thank Dr. Michael Tassia for helpful insights and Dr. Mercedes Burns, Prashant Sharma, Rajiv McCoy, and Erik Andersen for their helpful feedback and suggestions on the manuscript. This work was supported by …

## Data availability

Scripts, processed data and results can be found at github.com/GordusLab/orb-selection.

## Author Contributions

C.R. and A.G. conceived and designed the research strategy. J.M. compiled and pre-processed the transcriptomes. C.R. implemented all analyses. C.R.. and A.G. analyzed the data and prepared the manuscript.

---

**Supplementary Figure 1:** GO terms enriched in the sets of genes that showed significantly greater odds of having been lost among orb-weavers (yellow), duplicated among orb-weavers (green), lost among non-orb-weavers (blue), or duplicated among non-orb-weavers (red).

**Supplementary Figure 2:** Histograms of all 10,000 means and standard deviations of each of the permulated log odds ratio distributions of **(a)** gene loss and **(b)** gene duplication between orb-weavers and non-orb-weavers. Mean values of these statistics were used to generate average permulated distributions and confidence thresholds in Figure 5.

**Supplementary Figure 3:** Histograms of triple Gaussian parameters fit to each of the 10,000 permulated log odds ratio distributions are shown along with means of these permulated parameters and parameters of a triple Gaussian fit to the true distribution of log odds ratios of **(a)** gene loss and **(b)** gene duplication between orb-weavers and non-orb-weavers.

## Notes

### Competing Interest Statement

The authors have declared no competing interest.

