## Supplementary figures and images for "Comparative transcriptomic analysis reveals signatures of selection for orb-weaving behavior in spiders"

### Supplementary Fig. S1

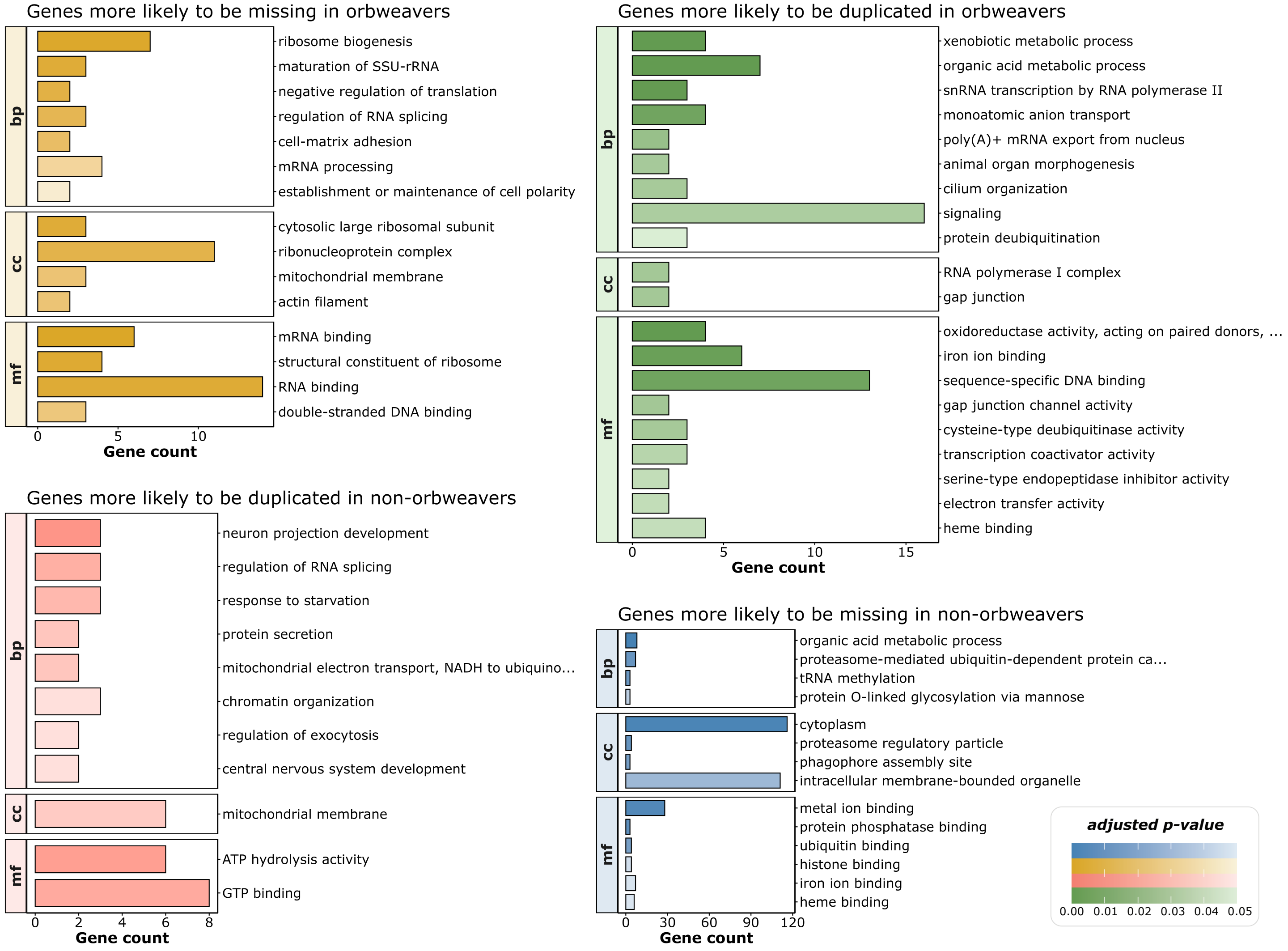

### Supplementary Fig. S2

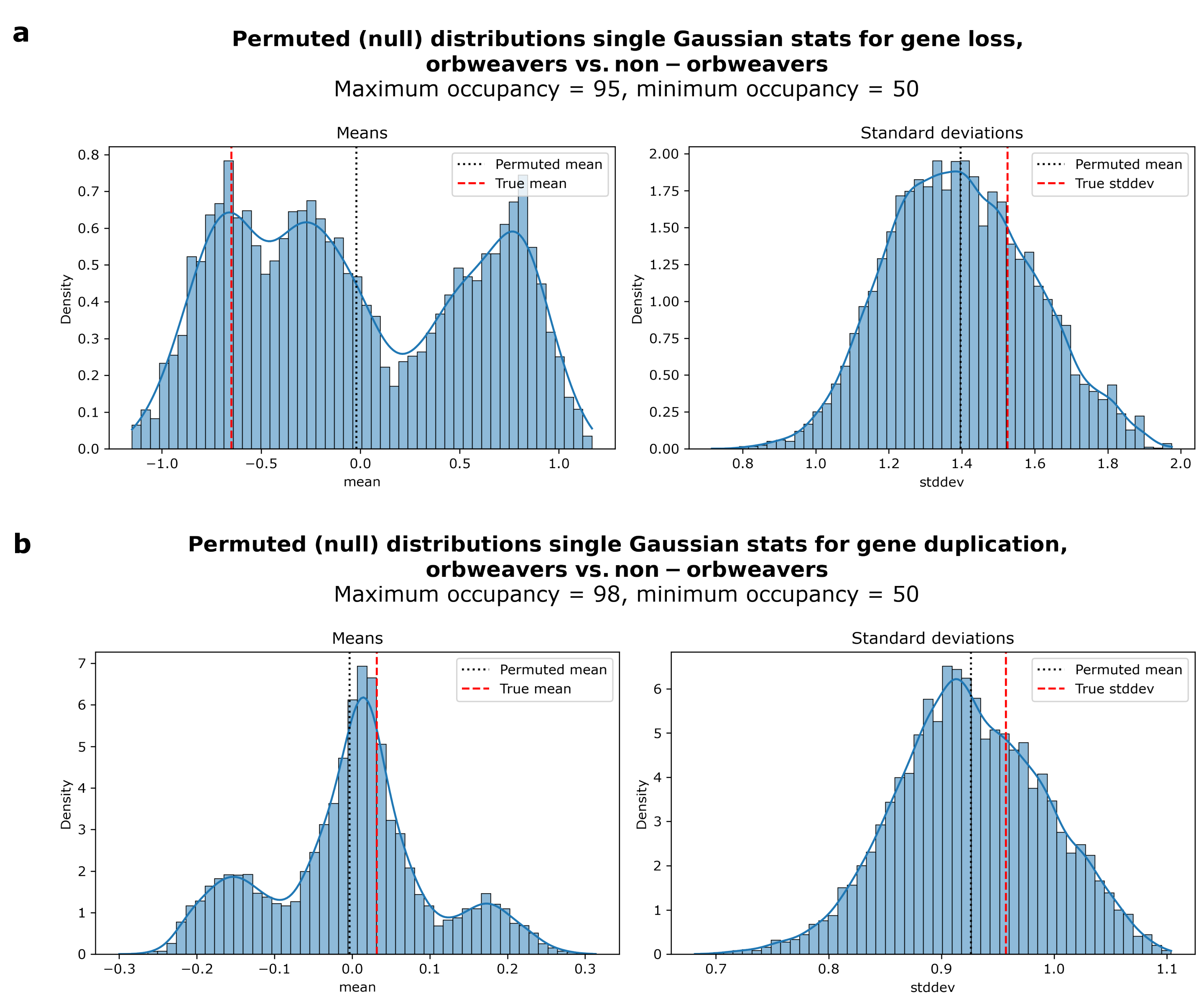

### Supplementary Fig. S3

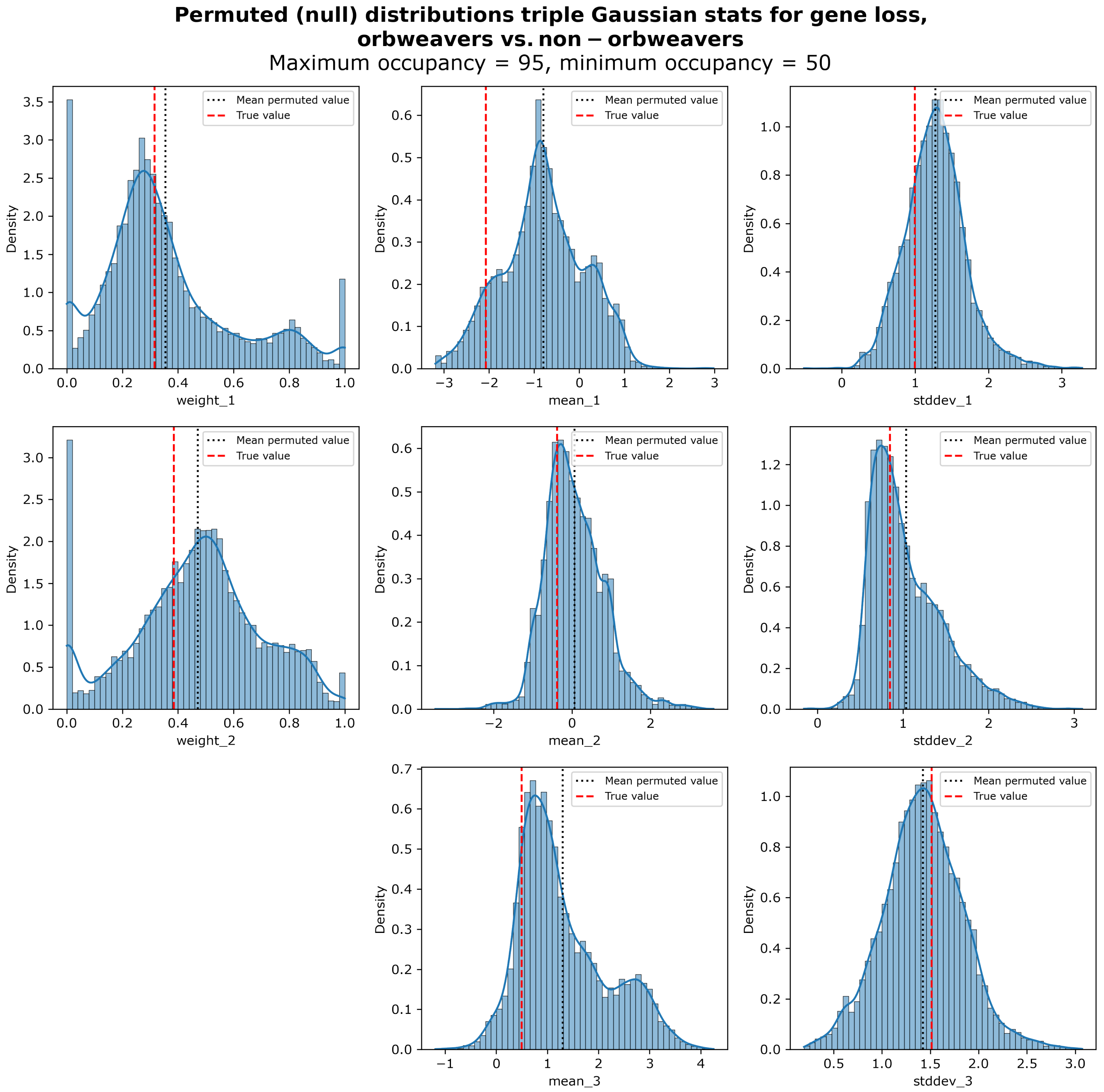

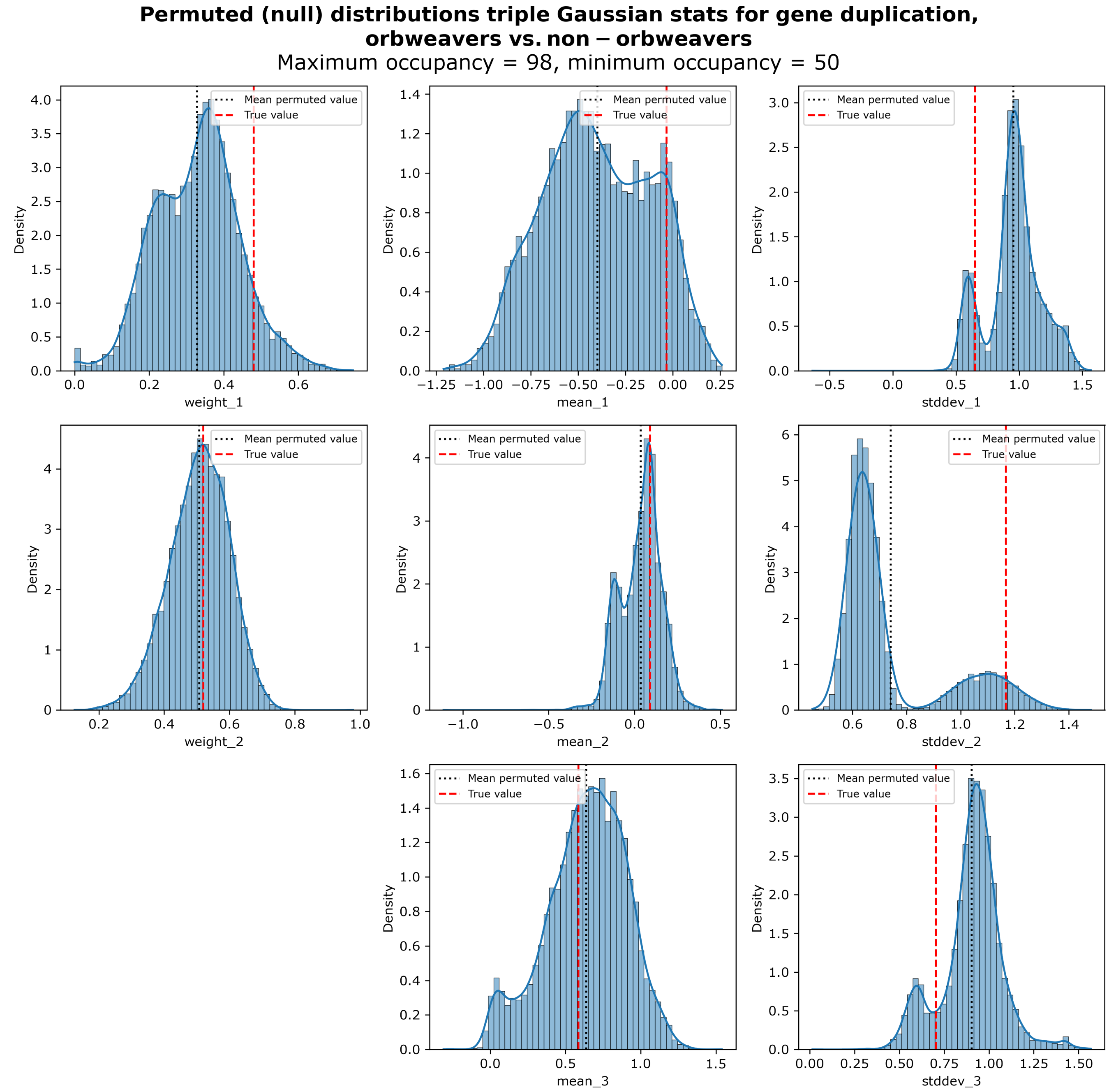
